# The Identification of Source and Vector of a Prolific Marine Invader

**DOI:** 10.1101/083972

**Authors:** Stacy A. Krueger-Hadfield, Nicole M. Kollars, Allan E. Strand, James E. Byers, Sarah J. Shainker, Ryuta Terada, Thomas W. Greig, Mareike Hammann, David C. Murray, Florian Weinberger, Erik E. Sotka

## Abstract

The source and vector of an introduced species inform its ecological and evolutionary history and may guide management that seeks to prevent future introductions. Surprisingly, few studies have successfully used genetic tools to independently inform the specific source and pathway of biological invasions. The ecological history of many introduced species, including their origins and vectors, is often based on suppositions or educated guesses. Here, we used mitochondrial and microsatellite genotyping to trace the invasion of the Asian seaweed *Gracilaria vermiculophylla* (Rhodophyta) along the three coastlines of the Northern Hemisphere to which it has been introduced: the western coast of North America, eastern coast of the United States and the coasts of Europe and northwest Africa. Analyzing 37 native and 53 introduced sites, we identified the Pacific coastline of northeastern Japan as the ultimate source of the Northern Hemisphere invasion. Coincidentally, most exports of the oyster *Crassostrea gigas* historically originated from this region and both species often grow in close proximity. Based on genetic signatures, each of the three coastlines likely received thalli directly from Japan, as well as material from another introduced coastline (i.e., a secondary invasion). Our ability to document a source region, which was enabled by a robust sampling of locations and loci that previous studies lacked, reflected strong phylogeographic structure along native coastlines. We suggest *Gracilaria vermiculophylla* is an important representative example of many species likely exported out of Japan by the oyster trade and its genetic signatures that may be a hallmark of oyster introduction legacies.

## INTRODUCTION

Non-native species represent one of the greatest threats to native biodiversity by homogenizing the Earth’s biota and altering community processes and ecosystem function. Predicting the environmental and evolutionary processes that facilitate invasions is one of the major goals of invasion biology (Kolar & Lodge 2001). For a species with a broad distribution and strong population genetic structure in its native range, the pooling of all native populations with which to compare against introduced populations may provide an uninformative contrast. Non-source populations in the native range may not share the same opportunities for invasion because the vector(s) may be absent (Miller & Ruiz 2008) or the necessary ecological or evolutionary characteristics that enable successful colonization are missing (i.e., a capacity for uniparental reproduction, e.g., Pannell *et al.* 2015). Comparisons that isolate the specific source region in the native range from which the introduced populations emanated can far better inform genetic, ecological and evolutionary patterns of invasion (Miller & Ruiz 2008). Therefore, elucidating invasion pathways is a fundamental and critical step to define and test hypotheses concerning the environmental and evolutionary processes underlying successful invasions (reviewed in Estoup & Guillemaud 2010). This knowledge will enable (i) an understanding of subsequent evolution in recipient regions (Bock *et al.* 2015), (ii) the identification of other invaders from the same region or as a result of the same vector (Brawley *et al.* 2009) and (iii) the development of effective management strategies (Dlugosch & Parker 2008).

For most invasions, our knowledge about pathways and vectors is largely based on historical and observational data that are often incomplete and misleading (Estoup & Guillemaud 2010). Population genetic tools and more advanced statistical analyses have facilitated our understanding of invasion pathways. In the marine environment, maritime activities, including shipping and aquaculture, have long been implicated in the introduction of non-native species (e.g., Ruiz *et al.* 2000; Grosholz 2002), particularly in estuarine habitats where these activities are more intense (reviewed in Ruiz *et al.* 1997; Preisler *et al.* 2009). Ballast water and hull fouling have received a great deal of attention (see for example Carlton & Geller 1993) and are often cited as the most likely vectors for many marine invasions (Ruiz *et al.* 2000). For example, shipping from the Gulf of Mexico and the eastern seaboard of the United States was the likely vector in the introductions of the ctenophore *Mnemiopsis leidyi* to the Black, Caspian, North, Baltic and Mediterannean Seas (Reusch *et al.* 2010). Voisin *et al.* (2005) found maritime traffic promoted recurrent invasions of the kelp *Undaria pinnatifida*, a species native to the northwest Pacific, to Australasia. Nevertheless, there is a surprisingly small subset of studies that include sufficient native and introduced regional sampling with which to document invasion histories and vectors in the marine realm.

The lack of robust sampling throughout native and introduced distributions coupled with a lack of appropriate population genetic tools hampers our understanding of marine invasions, particularly the underlying ecological and evolutionary processes. Moreover, the dearth of detailed invasion histories results in ambiguities about the vectors of ubiquitous invaders that are often characterized as polyvectic (see, for example, Grosholz *et al.* 2015). Particularly for organisms without a planktonic larval stage, aquaculture imports are a more likely vector (Grosholz *et al.* 2015). The Japanese oyster *Crassostrea gigas* is one of the main vectors facilitating the hitchhiking of numerous marine taxa throughout the near-shore marine environments in every ocean basin (Ruesink *et al.* 2005). Few studies, however, have provided convincing evidence linking the importation of *C. gigas* to the introduction of other nonindigenous species beyond coincidental occurrence in estuaries and anecdotes, though exceptions include Bonnot (1935) who documented the occurrence of invasive mollusks with imported Japanese oyster shipments. To our knowledge, Miura *et al.* (2006) is the only study that used molecular markers to re-trace an invasion to Northeastern Japan. Based on genotypic diversity in Japan and the western coast of North America, the invasion of the Asian mud snail *Batillaria attramentaria* invasion as well as one of its trematode parasites originated in the main region of Japanese *C. gigas* exports (Miura *et al.* 2006).

The haploid-diploid macroalga *Gracilaria vermiculophylla* (Ohmi) Papenfuss (Rhodophyta) has spread from its native distribution in the northwestern Pacific Ocean to virtually every high-salinity, temperate estuary in Europe and North America (Weinberger *et al.* 2008; Saunders 2009; Byers *et al.* 2012). *G. vermiculophylla* can profoundly transform estuarine ecosystems (e.g., Byers *et al.* 2012; Thomsen *et al.* 2013) and result in negative economic impacts (e.g., Freshwater *et al.* 2006). *G. vermiculophylla* was first identified through molecular barcoding from samples collected in 1979 in Baja California (Bellorin *et al.* 2004), 1994 in Elkhorn Slough (Bellorin *et al.* 2004), 1998 in the Chesapeake Bay (Thomsen *et al.* 2006) and 1995 in France (Rueness 2005). As a result, though studies have posited oysters as a vector (e.g., Rueness 2005; Thomsen *et al.* 2006), reviews of marine invasions suggest oysters are unlikely to be the main vector as the timing of oyster imports and the identification of *G. vermiculophylla* do not align. However, estuarine ecology does not have a strong phycological tradition as most macroalgae require abundant hard susbstratum with which to attach (Krueger-Hadfield *et al.* 2016) and this may have prevented early detection of *G. vermiculophylla* (Krueger-Hadfield *et al. in review*). Compounded by poor taxonomic resolution in the Gracilariales (Gurgel & Fredericq 2004), cryptic invasions were likely occurring earlier in North America and Europe based on herbarium surveys (SA Krueger-Hadfield and KA Miller, *unpubl. data*). Thus, the vectors that facilitated the movement of *G. vermiculophylla* thalli across oceanic basins remain speculative.

Despite a flurry of studies assessing the impacts of the invasion (e.g., Nyberg & Wallentinus 2009; Thomsen *et al.* 2013; Hammann *et al.* 2013b), only a single investigation has attempted to trace the invasion of *Gracilaria vermiculophylla* throughout the Northern Hemisphere. Kim *et al.* (2010) suggested the Sea of Japan/East Sea was the source of the invasion. Yet, their study included only five Japanese sites, none of which occurred on the northeastern coastlines of Honshu and Hokkaido, areas of important aquaculture activities not only for *C. gigas* (Byers 1999), but also agar extraction from *Gracilaria* spp., including *G. vermiculophylla* (Okazaki 1971). Due to the uncertainty of the invasion source(s), the investigation of hypotheses that test the underlying evolutionary processes facilitating this widespread invasion is limited.

Previously, we sampled 30 native sites, with an emphasis on Japanese sites (23 of the 30 sites), and 35 introduced sites along the coastlines of western and eastern North America and Europe (Krueger-Hadfield *et al.* 2016). We found comparable levels of genetic diversity between native and introduced sites, suggesting highly genetically diverse incocula or multiple invasions (Krueger-Hadfield *et al.* 2016). Introduced sites, however, generally had a lack of abundant hard substratum, a necessity for algal spore recruitment. As a result, introduced sites were dominated by diploid tetrasporophytes (> 80% diploid thalli, on average) and showed genetic signatures of extensive vegetative fragmentation (Krueger-Hadfield *et al.* 2016). Uniparental reproduction, such as clonality, is one of the main attributes of successful colonizers, including invaders (Kolar & Lodge 2001; Pannell *et al.* 2015), and likely facilitated the widespread invasion of *G. vermiculophylla*.

Here, we focus on spatial genetic structure of the samples previously described (Krueger-Hadfield *et al.* 2016) as well as adding 25 new native and introduced sites to investigate the invasion history of *G. vermiculophylla*. First, we explored the patterns of genetic structure and differentiation in mitochondrial and microsatellite loci along three native coastlines: China, South Korea and Japan. Second, we tested the hypothesis that the northeastern coast of Japan, particularly sites in the Miyagi and Hokkaido Prefectures, served as the source of the Northern Hemisphere invasion. Third, we investigated the patterns of genetic structure and differentiation within and among sites along each of the introduced coastlines in order to determine the number of invasions and the occurrence of secondary spread.

## MATERIALS AND METHODS

### Data generation

#### Sample collection

Algal thalli were sampled from 90 sites across the Northern Hemisphere from 2012-2016 (Table S1). Thirty-seven sites were from the native range and sampled along the coastlines of China, South Korea and Japan (Figure 1, Table S1, Figure S1). Fifty-three sites were sampled along the coastlines of western North America (hereafter, WNA), the eastern United States (hereafter, EUSA) and northern Africa and Europe, including the British Isles (hereafter, EU; Figure 1, Table S1, Figure S1). Of these 90 sites, 30 native and 35 introduced sites were used in a previous study in which we compared the reproductive system between native and introduced regions (Krueger-Hadfield *et al.* 2016).

**Figure 1.**
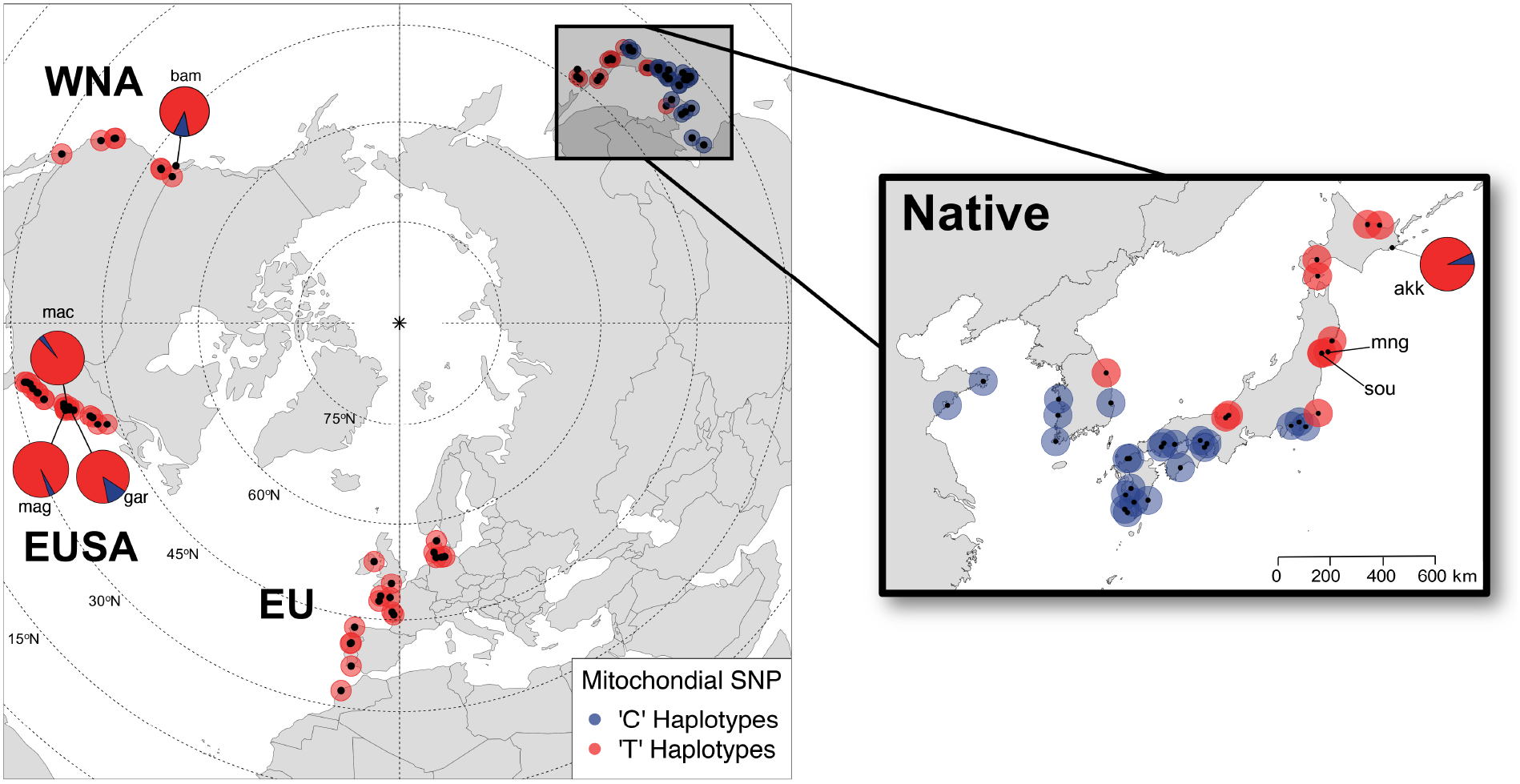
Geographic distribution of the mitochondrial SNP frequency among sampled sites of *Gracilaria vermiculophylla* in the native range (Northwest Pacific) and non-native range (WNA = western North America; EUSA = eastern United States; EU = British Isles, continental Europe and Northern Africa). At most sites, thalli were either ‘C’ (blue) or ‘T’ (red) at basepair 945 of the sequenced *cox*1 gene from Kim *et al.* (2010). At five sites, both ‘C’ and ‘T’ SNPs were detected and relative frequencies are shown by pie charts. More than half of the introduced thalli were assigned to one of three native sites (akk, mng or sou) using DAPC (see also Figure S6) and are shown in the native map. Sampling information is provided in Table S1 and mitochondrial sequencing and genotyping data are provided in Figure S1 and Table S2.

The species identity of all thalli was confirmed either using DNA barcoding (Kim *et al.* 2010), the amplification of 10 species-specific markers (Kollars *et al.* 2015; Krueger-Hadfield *et al.* 2016) or both. *Gracilaria vermiculophylla* thalli from native sites were attached via holdfasts to hard substrata (i.e., bedrock, pebbles, gastropod shells), whereas, in the introduced range, there was a spectrum of sites ranging from fully attached to fully free-floating *G. vermiculophylla* thalli (Krueger-Hadfield *et al.* 2016, Krueger-Hadfield and Sotka, *unpubl. data*). At all sites, samples were collected with at least one meter separating each putative genet to avoid sampling the same genet twice (Guillemin *et al.* 2008; Krueger-Hadfield *et al.* 2016). Ploidy was determined by observing reproductive material under a binocular microscope (40x) and/or using microsatellite loci (Krueger-Hadfield *et al.* 2016).

#### DNA extraction, microsatellite and mitochondrial cox1 amplification

Total genomic DNA was isolated and microsatellite simplex PCRs and fragment analysis were performed following Krueger-Hadfield *et al.* (2016; see also Table S1). However, for sites sampled in 2015-2016 (Table S1), we only extracted phenotypically diploid thalli as part of a larger study on adaptation among native and introduced regions (Sotka *et al. in review*). In total, we genotyped 2192 diploid and 785 haploid thalli across 90 native and introduced sites.

The mitochondrial gene *cox*1 was amplified using the primer sets 43F (Geraldino *et al.* 2006) and 880R (Yang *et al.* 2008) and 622F (Yang *et al.* 2008) and 1549R (Geraldino *et al.* 2006) for 201 thalli sampled across both the native and introduced ranges (Table S2, Figure S2). We did not distinguish between haploid and diploid thalli for the mitochondrial sequencing. PCR amplification was performed on a total volume of 25 μL, containing 0.5 U of *taq* DNA polymerase, 2.5 mM of each dNTP, 2 mM MgCl_2_, 1 x reaction buffer, 250 nM of each primer and 5 μL of DNA and PCR conditions previously described (Yang *et al.* 2008). Approximately, 5 μL of PCR product with 1 μL of Orange G loading dye were visualized on 1.5% agarose gels and 1 μL of ethidium bromide.

One μL of ExoSAP-It was added to 7 μL of PCR product (Affymetrix, Santa Clara, CA, USA) and incubated for 15 minutes at 37 °C followed by 15 minutes at 80 °C. Four microliters of 2 μM primer was added to each product and sequenced in the forward direction commercially by Eurofins Genomics (Louisville, KY, USA). Sequences were edited using *4Peaks* (Nucleobytes, The Netherlands), aligned with the haplotypes from Kim *et al.* (2010) using *Muscle* (Edgar 2004) in *Seaview* ver. 4.6 (Gouy *et al.* 2010) with default parameters. We assigned haplotype numbers using Kim *et al.* (2010), which sequenced a ˜1200 base pair fragment. We also defined six new ˜1200 base pair haplotypes that were not previously sequenced (Kim *et al.* 2010) using *DnaSP*, ver. 5.10.1 (Librado & Rozas 2009). We ignored the haplotype designations of Gulbransen *et al.* (2012) because their haplotypes covered only a subset of this larger ˜1200 base pair fragment.

#### RFLP assay and data analysis of mitochondrial sequences

We discovered a single nucleotide polymorphism (C/T) at the 945^th^ base pair within the mitochondrial *cox*I fragment which delineated among several haplotypes. Furthermore, the restriction enzyme Af1III (New England Biolabs, Ipswich, MA, USA) could distinguish the alleles at this SNP (ACGTG(T/C; Table 1, Table S2, Figure S3). Restriction enzyme digestion was performed on thalli from each site using a total volume of 25 μL, containing 1X buffer, 10 U of AF1III and 5 μL of the *cox*1 PCR product amplified using the primers 622F (Yang *et al.* 2008) and 1549R (Geraldino *et al.* 2006) and under the following reaction conditions: 37 °C for 2 hours, 80 °C for 20 minutes and 20 ^0^C for 5 minutes. Restriction digest products were visualized on 1.5% agarose gels with 1 μL of ethidium bromide (Figure S1c).

**Table 1.** Haplotypic diversity across the native and introduced ranges of *Graclaria vermiculophylla*. a) Sites south and north of approximately 35°N in China, South Korea and Japan were delineated based on C:T frequencies (see also Figure 1). We used the haplotype numbers assigned by Kim *et al.* (2010) as these covered the ˜1500 bp of *cox*1. We have extended Kim *et al.* (2010) and generated haplotype numbers for six new haplotypes (Haplotypes 26 – 31). The proportion of the ‘C’ and ‘T’ SNPs are given for each region (see Table S1 for sampling sizes are each site). b) The biogeographic province {Briggs:2011cy} and ecoregions {Spalding:2007th} in which each of the native range haplotypes were found.

| Region | ‘C’ Haplotypes | ‘T’ Haplotypes | C:T Ratio |
| --- | --- | --- | --- |
| South of ~35°N | 1 – 4, 8 – 15, 17, 27, 30 – 31 | None | 1.00 : 0.00 |
| North of ~35°N | 26 | 5 – 7, 16, 18, 28 | 0.01 : 0.99 |
| WNA | 15 | 6, 19, 29 | 0.02 : 0.98 |
| EUSA | 15 | 6 | 0.01 : 0.99 |
| EU | None | 6, 18 | 0.00 : 1.00 |

b)
| Biogeographic province | Ecoregion | ‘C’ Haplotypes | ‘T’ Haplotypes |
| --- | --- | --- | --- |
| Cold temperate - Kurile | Sea of Okhotsk | None | 6 |
|  | Oyashio Current | 26 | 6 |
|  | Northeastern Honshu | None | 6 |
| Cold temperate - Oriental | Sea of Japan | None | 5 – 7 |
|  | Northeastern Honshu | None | 6, 18, 28 |
|  | Yellow Sea | 1 – 2, 17 | None |
| Warm temperate – Sino-Japanese | Sea of Japan | 4 | None |
|  | East China Sea | 2 – 4; 8 – 13; 30<br>– 31 | None |
|  | Central Kuroshio Current | 14 – 15; 27 | 16 |

A phylogeny of the haplotypes from Kim *et al.* (2010) and new haplotypes uncovered in this study was constructed using the Hasegawa-Kishino-Yano model (HKY) plus gamma (Hasegawa *et al.* 1985), one of the appropriate models detected using ModelTest (Guindon & Gascuel 2003, Darriba *et al.* 2012), as implemented in MEGA6 (Tamura *et al.* 2013). The resulting trees were edited using *FigTree* ver. 1.4.2 (http://tree.bio.ed.ac.uk/software/figtree/).

### Data analysis

#### Ploidy and genet determination

To analyze both native and introduced ranges, only the diploid thalli were used for subsequent microsatellite data analyses (see also Krueger-Hadfield *et al.* 2016).

Custom *R,* ver. 3.2.2 (R Core Team 2016) routines were written in order to determine ploidy based on the multilocus genotype and create input files for downstream analyses. The number of repeated identical multilocus microsatellite genotypes (MLGs) was computed using *genclone* ver. 2.0 (Arnaud-Haond & Belkhir 2006) and a custom *R* routine following (Parks & Werth 1993; Arnaud-Haond *et al.* 2007). *P_sex_*, or the probability for a given MLG to be observed in *N* samples as a consequence of different sexual reproductive events, was calculated for each repeated MLG. If *P_sex_* was greater than 0.05, duplicated multilocus genotypes were considered as different genets. If *P_sex_* was smaller than 0.05, the duplicated MLGs were considered as ramets (or clones) of the same genet.

Studies on the population genetics of clonal organisms remove repeated MLGs before calculating heterozygosity and other *F*-statistics in order to avoid distorting these estimates (e.g., Sunnucks *et al.* 1997; Halkett *et al.* 2005; Krueger-Hadfield *et al.* 2011). Following this convention, we chose the conservative approach of performing the following analyses for diploid MLGs for which *P_sex_* > 0.05 (i.e., one-thallus-per-genotype based on *P_sex_*).

#### General description of genetic variation

For each site, the average expected heterozygosity (*H_E_*) was calculated using *GenAlEx*, ver. 6.5 (Peakall & Smouse 2006; Peakall & Smouse 2012). An estimate of the mean expected number of alleles (*A_E_*) was computed using rarefaction implemented in the program *HP-Rare,* ver. 1.0 (Kalinowski 2005) on the smallest sample size of 10 diploids (i.e., 20 alleles). Finally, the number of expected MLGs (*eMLGs*) at the smallest sample size (n = 10 diploid thalli) was estimated using rarefaction in the *R* package *poppr,* ver. 2.2.1 (Kamvar *et al.* 2014; 2015). For the latter two diversity metrics, sites with less than 10 diploid thalli were excluded from calculations.

We calculated pairwise genetic differentiation based on allele identity (*F_ST_*) and allele size (*ρ_ST_*) among sites along each of four coastlines: Japan, WNA, EUSA and EU using *genpop* ver. 4.4 (Rousset 2008). We measured geographic distance (km) following the contours along coastlines among all site pairs using the measure distance function in Googleβ^®^ Maps. We performed Mantel tests in order to detect relationships between the genetic and geographic distance along each of the four coastlines using *R* (R Core Team 2016).

#### Multivariate analyses of microsatellite data

To assess relationships among multilocus genotypes, we pursued multivariate analyses that avoided making strong assumptions about the underlying genetic model (Jombart *et al.* 2009). Discriminant analysis of principal components (DAPC) is one such class of analyses, in which principal components that best summarize the differences among clusters are found, while also minimizing within-cluster variation (Jombart *et al.* 2010). This technique extracts information from genetic datasets by performing a principle component analysis (PCA) on pre-defined groups or populations (see below). We performed DAPC using the package *adegenet,* ver. 2.0.1 (Jombart 2008; Jombart & Ahmed 2011) implemented in *R* ver. 3.2.2 (R Core Team 2016). The PCA factors were used as variables for a discriminant analysis that maximizes the inter-group component of variation. The PCA ensures the variables using in the discriminant analysis are uncorrelated, thus removing any potential effects of linkage disequilibrium.

We performed the DAPC with increasing numbers of PCs on 90% of our data and then the remaining 10% of the individuals were projected onto the discriminant axes constructed by the DAPC. It was possible to measure how accurately the remaining 10% of the individuals were placed in multidimensional space (i.e., how well their position corresponds to their group membership). Based on this cross-validation with the *xvalDapc* function, we retained 88 principal components (PC) that explained 87.8% of the total variance for subsequent DAPC’s (Figure S2a).

We estimated how well-supported the group membership was relative to five *a priori* subregions using the *compoplot* function in *adegenet*. Posterior group memberships were utilized in order to indicate admixture or the misclassification when prior groups are used to conduct the DAPC. Based on the mitochondrial SNP, we divided the native range into the ‘C’ haplotype group and the ‘T’ haplotype group corresponding to the break between 35°N and 37.5°N in Japan and South Korea in which the two SNPs dominated to the south and north, respectively (Figure 1, Table 1). For the introduced range, we treated each coastline as a different group: WNA, EUSA and EU. The stability of *a priori* group membership probabilities, derived from proportions of successful reassignments based on retained discriminant functions of DAPC, were high (> 92%) for the native ‘C’ haplotype, EUSA and EU subregion, but lower for the native ‘T’ haplotype and WNA subregions (85.3% and 76.4%, respectively; Figure S2). Based on these reassignment frequencies, we performed the DAPC with the aforementioned five subregions.

Next, we predicted the assignment of thalli from one region to another using supplementary individuals that were not used in model construction as implemented in *adegenet*. We performed this analysis with training data composed of (i) only the native sites and (ii) sites from the native range and two of the three introduced coastlines. We transformed ‘new data’ using the centering and scaling of training data (i.e., (i) the native range alone or (ii) the native range and the two introduced coastlines). Using the same discriminant coefficients, we predicted the position of the new individuals (i.e., (i) all introduced sites and (ii) the one introduced coastline not used in the training data).

#### Bayesian clustering of microsatellite data

Because both native and introduced populations of *Gracilaria vermiculophylla* are out of Hardy-Weinberg equilibrium (Krueger-Hadfield *et al.* 2016), we chose to use Bayesian model-based clustering that relaxes the HWE assumption as implemented in the software *instruct* (Gao *et al.* 2007). Simulations were performed using *instruct* with a model including both bi-parental inbreeding and admixture, where each individual drew some fraction of its genome from each of the *K* populations. A burn-in of 300,000 repetitions and a run length of 500,000 were used for *K* = 2 to *K* = 30, where 20 iterations for each *K*.

To evaluate the values of *K*, we analyzed *K* = 2 to 30 clusters using *clumpak* (Kopelman *et al.* 2015). *clumpak* identifies sets of highly similar runs across the 20 independent runs of each *K* generated with *instruct* and separates them into distinct major and minor modes. *clumpak* utilizes the software *clumpp* (Jakobsson & Rosenberg 2007) in order to generate a consensus solution for each distinct mode using a Markov clustering algorithm that relies on a similarity matrix between replicate runs. Next, *clumpak* identifies an optimal alignment of inferred clusters and matches the clusters across the values of *K* tested.

We determined the optimal number of *K* using outputs from *instruct* and *clumpak*. First, we plotted the number of clusters against the values of DIC (+ SE) and chose the value of *K* at the point at which the curve reached an asymptote. The lower the DIC value, the better fit of the model used (i.e., the number of K). Second, we used the number of independent runs out of 20 that generated the major mode and the mean similarity score from *clumpak.*

#### Clustering analyses of microsatellite data

We constructed a neighbor-joining (NJ) tree based on Jaccard distance using the *R* packages *ape* ver. 3.5 (Paradis *et al.* 2004) and *phytools* ver. 0.5-38 (Revell 2011). The resulting trees were edited using *FigTree* ver. 1.4.2 (http://tree.bio.ed.ac.uk/software/figtree/).

## RESULTS

### Mitochondrial structure across native and introduced ranges

Across 201 thalli sequenced from the native and introduced ranges, we identified 10 distinct *cox*1 haplotypes (Figure S1, Table S2), four of which were described previously by Kim *et al.* (2010). Three clusters of haplotypes were identified with a maximum-likelihood (ML) analysis based on 19 previously described haplotypes (Kim *et al.* 2010) and the six new haplotypes from this study. The three clades were heterogeneously distributed across the native range (Figure S1a). Two clades were found south of ˜35 °N latitude: one consisted of 10 haplotypes including haplotypes 1, 2, 3, 9, 10, 11, 12, 17, 26, 30, 31 and the other clade of six haplotypes included 4, 8, 13, 14, 15 and 27 (Table 1, Figure S1a). The third clade of six haplotypes were only found sequenced north of ˜35 °N in the native range (Figure 1, Table 1, Figure S1). Seventy-two of 77 thalli, or 94% of sequenced thalli, collected north of 35°N were haplotype 6, the haplotype that is known to dominate the introduced range (Kim *et al.* 2010).

Two single nucleotide polymorphisms, or SNPs (basepair 945 and 1040 from Kim *et al.* 2010, Table S2) delineated all haplotypes that occur north (‘T’) from those that occurred south (‘C’) of ˜35°N in the native range (Figure S1, Table S2). Based on RFLP genotyping of 691 thalli collected south of ˜35 °N latitude, we found that all were ‘C’ haplotypes (Figure 1, Table 1). An RFLP analysis of 375 thalli collected north of ˜35°N revealed all were ‘T’ haplotypes. The exception was three thalli from Akkeshi (akk) that were ‘C’ belonging to the new haplotype 26 that diverges from all other haplotypes by at least 5 base pairs (Figure S1a).

In the introduced range, 99.5% (1597 of 1605 thalli) of the thalli were ‘T’ haplotypes and were dominated by haplotype 6 (82 of 101, or 81%, sequenced thalli; Figure 1, Table 1, Figure S1b-d, Table S2). Only nine introduced thalli were ‘C’ haplotypes (3 from Bamfield/bam, 4 from Gargatha/gar, 1 from Machipongo/mac and 1 from Magotha/mag) and all belonged to haplotype 15. Haplotype 15 was originally sampled at Funabashi in Tokyo Bay (Kim *et al.* 2010), near the ‘C/T’ haplotype break. The ‘T’ haplotypes, 19 (Elkhorn Slough/elk; Kim *et al.* 2010) and 29 (Bamfield/bam; this study) were detected in the introduced range, but not in the native range, likely because of under-sampling in northern Japan (Table S2). For example, the ‘T’ haplotype 18 was previously only sampled in France (Kim *et al.* 2010), but was found after sequencing thalli from Mangoku-ura (mng; Figure S1, Table S2).

Taken together, mitochondrial data suggest the region north of ˜35 °N latitude was the source of the widespread Northern Hemisphere invasion (Figure 1). However, we could not resolve the sites that contributed to the invasion at finer spatial scales due to the lack of mitochondrial polymorphism within this northern native region.

### Microsatellite structure in native and introduced ranges

Microsatellite loci revealed profound genetic structure within the native range and identified cryptic genetic structure among and within introduced coastlines that could not be detected with mitochondrial genotyping. We focus on each of these patterns in turn.

Within the native range, microsatellite genotypes distinguished northern sites dominated by ‘T’ *cox*1 haplotypes and southern sites dominated by ‘C’ haplotypes. This separation was evident along the 1^st^ axis of a multivariate DAPC, which itself explained 55% of overall variation (Figure 2). ‘T’ versus ‘C’ differentiation was also confirmed by Bayesian clustering analyses in which sites were placed into subregions corresponding to genetic similarity (Figure 3, Table S3). At *K*=5, the optimal *K* based on the similarity score using *clumpak* and the curve of DIC estimates, showed strong differentiation within and between the ‘T’ and ‘C’ regions in the native range (see Figure S3 b–f for individual-level analyses and across *K*’s).

**Figure 2.**
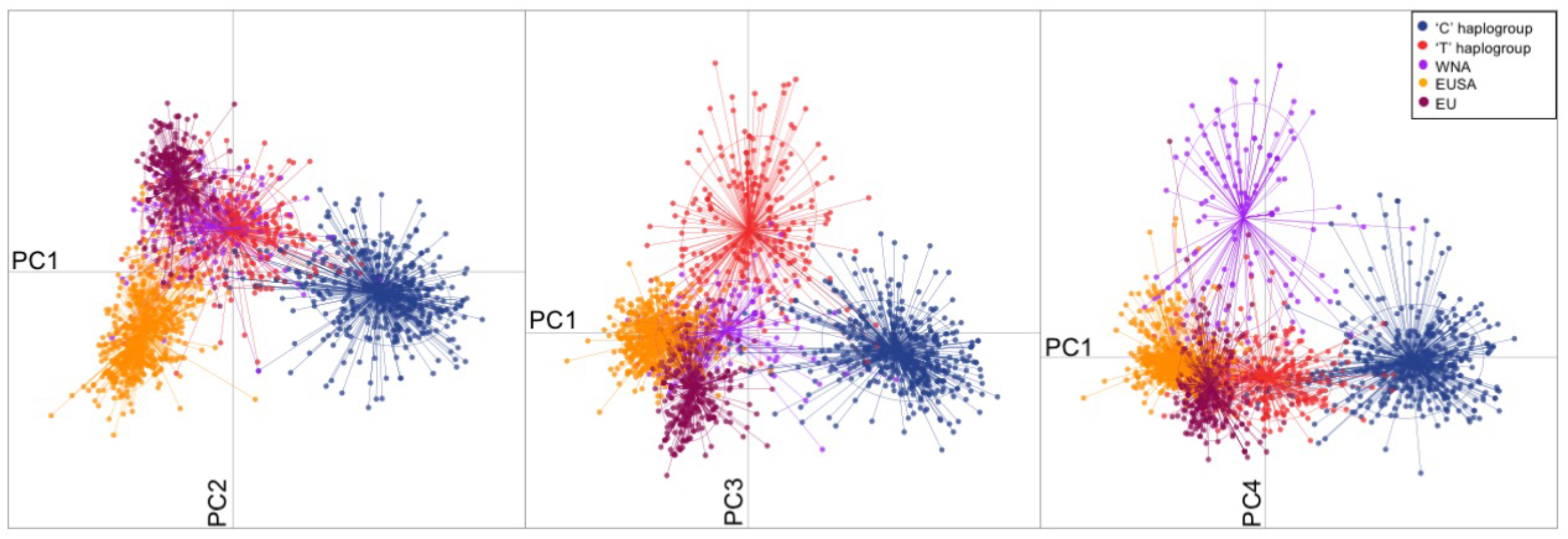
DAPC (Discriminant Analysis of Principal Components) relationships among microsatellite genotypes of *Gracilaria vermiculophylla*. We color-coded individuals corresponding to five *a priori* groups with high reassignment frequencies (see Figure S2) as the ‘C’ haplogroup (native sites dominated by the ‘C’ mtSNP and south of ˜35°N), the ‘T’ haplogroup (native sites dominated by the ‘T’ mtSNP and north of ˜35°N), WNA (western North America), EUSA (eastern United States of America) and EU (Europe and northern Africa). The first four principal components are shown (PC1: 55.4%, PC2: 19.4%, PC3: 14.3%, PC4: 10.9%).

**Figure 3.**
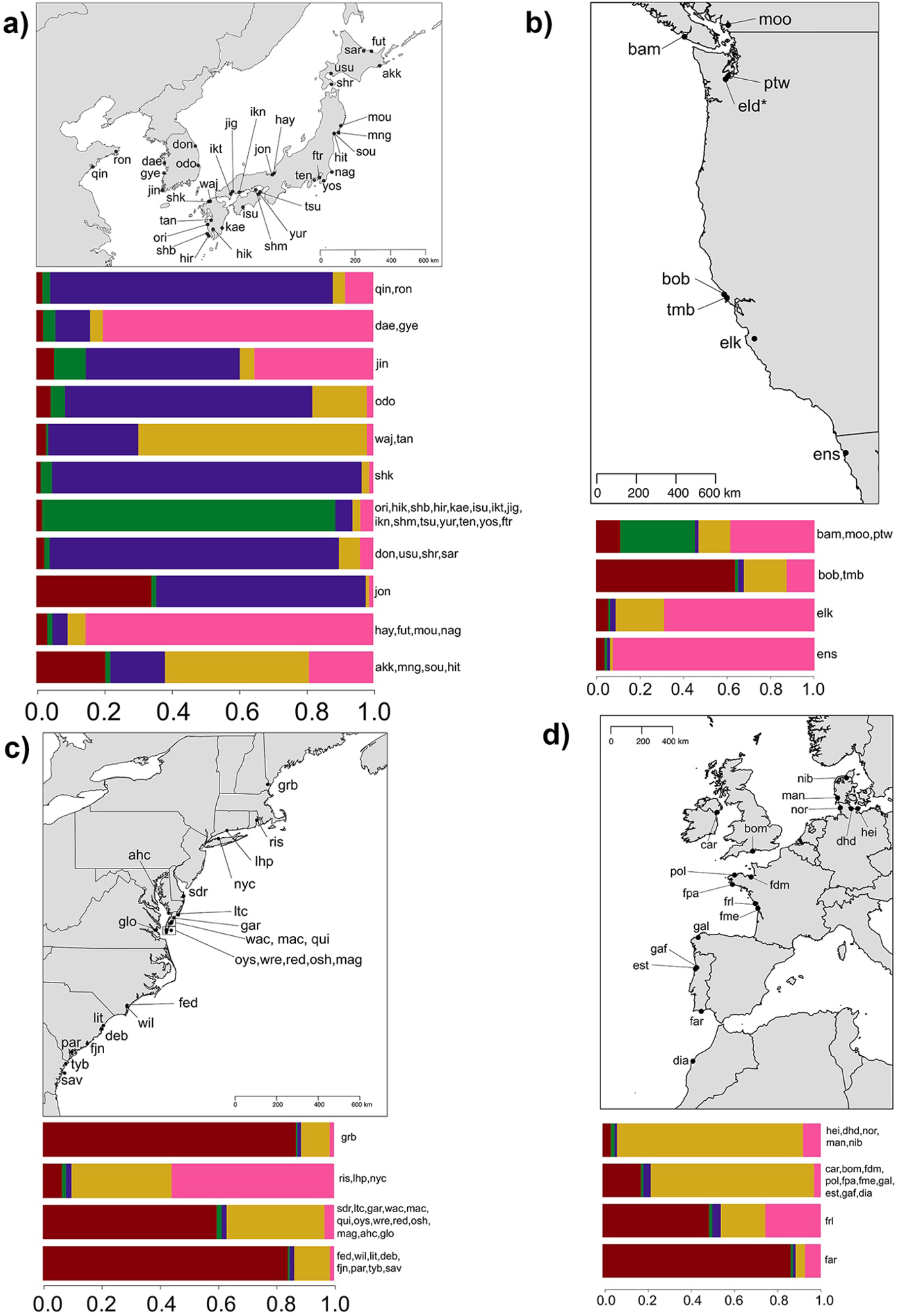
The mean assignment to five genetic clusters (colored as red, green, purple, gold and pink) as generated using *instruct* Gao *et al.* (2007) and *clumpak* Kopelman *et al.* (2015). We grouped thalli across sites of similar genetic composition using a visual inspection of individual cluster assignment (see Figure S3e) and used barplots to display individual assignments averaged across sites for that group. * As Eld Inlet had only one thallus based on *P_sex_*, it was excluded from Bayesian analyses, but is shown on the map.

At finer geographic scales, there was a high degree of genetic differentiation among sites that were close in proximity (i.e., less than 100 km) in the native range. Genetic differentiation in the native range, as measured using allele identity (*F_ST_*) and allele size (*ρ_ST_*), was high on average (*˜*0.47, Figure S4, Table S4). Genetic isolation was positively correlated with geographic distance when measured using allele identity (*F_st_*), but not with allele size (*ρ_ST_*, Figure S4, Table S4).

In the introduced range, the three continental shorelines were genetically differentiated and, thus, likely reflect different invasion histories. The DAPC analyses showed a separation of eastern United States (labeled as EUSA) thalli from the northern Japanese sites and other introduced shorelines along the 2^nd^ principal component axis, which itself explained ˜19% of the variation (Figure 2). The British Isles, European and northern African (labeled as EU) and western North American (labeled as WNA) thalli were differentiated from other regions along the 3^rd^ (14.3% of variation) and 4^th^ (10.9% of variation) principal component axes, respectively. With increasing numbers of genetic clusters using Bayesian analyses, introduced coastlines showed greater levels of genetic divergence from each other. At *K*=3, WNA sites were differentiated from the majority of EUSA and EU sites (Figure S3c), while at *K*=5, the EUSA and EU were genetically distinguishable (Figure S3e).

The strength of differentiation among populations along a shoreline differed across continental margins. Within WNA and EU shorelines, sites exhibited greater levels of genetic differentiation (average *F_ST_ ˜* 0.33 and *ρ_ST_* ˜ 0.20; Table S4) than was seen along the EUSA (average *F_ST_ ˜* 0.15 and *ρ_ST_* ˜ 0.18; Table S4). Significant patterns of isolation by distance (IBD) were detected along WNA and EU coastlines, but IBD was only marginally significant along the EUSA when measuring differentiation by allele identity (*F_ST_*; Figure S4, Table S4). In contrast, there were positive relationships between genetic differentiation and geographic distance along the EU and EUSA when measuring differentiation by allele size, but they were only marginally significant (*ρ_ST_*, Figure S4, Table S4).

It is likely that among-site differentiation within the introduced range reflected complex genetic origins that differ between continental margins. Along the WNA coastline, the Pacific Northwest (bam, pmo and ptw), Californian (bob and tmb), Elkhorn Slough (elk) and Ensenada (ens) were composed of different genetic clusters (Figure 3; Figure S3). Similarly, a neighbor-joining clustering analysis of site-level genetic distances indicated that the WNA coastline has been separately invaded on two or three occasions, given that sites are in separate locations across the tree (Figure S5). However, the bootstrap support for the NJ tree was poor and, as a result, this analysis should be interpreted with caution.

Along the EUSA coastline, most sites were composed of a homogenous set of genetic constituents, regardless the number of genetic clusters (Figure 3, Figure S3). The exceptions were the rocky-shore sites of Long Island Sound (lhp and nyc) and Narragansett Bay (ris), which seemed to have a genetic constituent more closely aligned with WNA and European populations (Figure 3, Figure S3). In the NJ clustering analysis, the majority of the EUSA sites were also part of the same clade, with the exception of these three northeastern sites (ris, lhp and nyc; Figure S5).

Along the EU coastline, Bayesian clustering revealed differences between sites sampled in Germany and Denmark (hei, dhd, nib, man, nor) versus a group of French (frl) and Portuguese populations (far), while the remaining populations in Ireland, the United Kingdom, France, Spain, Portugal and Morocco reflected admixture and a similar set of genetic constituents (Figure 3, Figure S3). The NJ tree suggested a largely similar origin of all European populations, with the exception of the site in Dorset (bom) in the United Kingdom which clustered with a site in the Chesapeake Bay (ahc; Figure S5).

There was a relationship between different measures of genetic and genotypic diversity (*H_E_, A_E_* and *eMLG*) and latitude along the EUSA and for genetic diversity metrics (*H_E_* and *A_E_*) along the WNA coastlines (Figure 4, Table S5). Highest diversity was found in the mid-latitudes along each coastline. Similar patterns were found along the coastlines of Europe and northern Africa, but none of the patterns were significant (Table S5).

**Figure 4.**
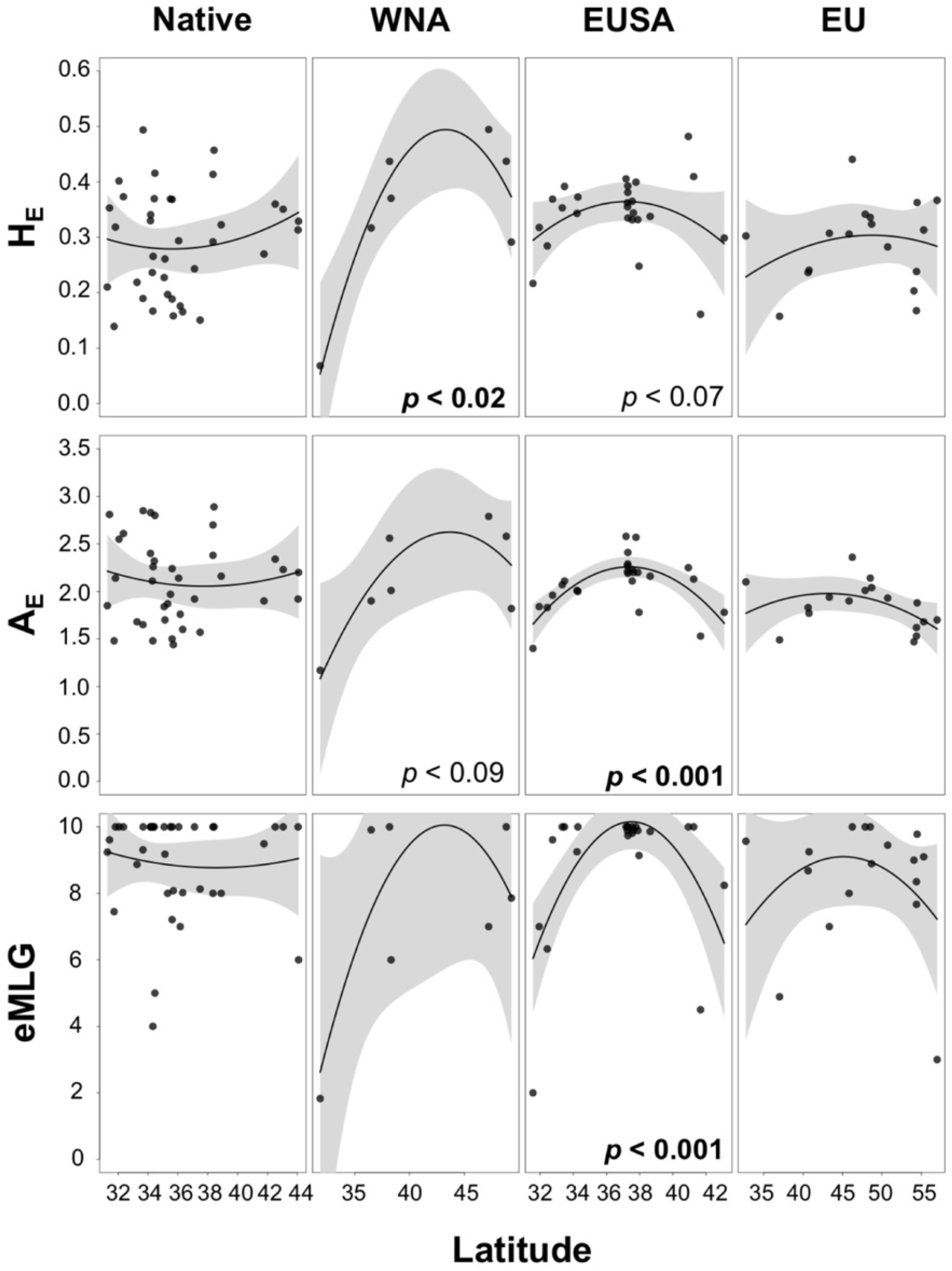
Genetic (*H_E_* and *A_E_*) and genotypic (*eMLG*) diversity as a function of latitude along the coastlines of the native range, WNA, EUSA and EU. Significance values are shown in the plots and bold if less than *p* = 0.05.

### Population sources for primary and secondary introductions

Several analyses of microsatellite data indicated the Pacific shoreline of northeastern Japan was the ultimate genetic source of introduced *Gracilaria vermiculophylla* throughout the Northern Hemisphere. Blind assignment of introduced populations onto multivariate DAPC space created by genotypes from native sites suggested most introduced thalli originated in three northern Japanese sites (791 thalli; ˜83%): Soukanzan (sou), Mangoku-ura (mng) and Akkeshi (akk, Figure 1, Figure S6). Of the remaining 164 thalli, 101 (˜11%) were assigned to other northern ‘T’ haplotype sites, principally along northeastern Honshu and eastern Hokkaido (Figure 1, Figure S6). Of the remaining 63 thalli, 14 WNA thalli (˜2%) were assigned to Yoshio (yos), a site on the Chiba Peninsula at the ‘C/T’ break, and 15 EU thalli (˜2%) and 22 EUSA thalli (˜2%) were assigned to Shikanoshima (shk), located in northern Kyushu (Figure 1, Figure S6).

Both Bayesian (Figure S3) and neighbor-joining clustering methods (Figure S5) supported the notion that introduced coastlines were most closely aligned with the Pacific shoreline of northeastern Japan. With increasing *K*’s, the genetic clusters found in the northern ‘T’ haplotype sites were the same clusters found predominately along each introduced coastline (Figure 3, Figure S3). Similarly, NJ analyses also confirmed genetic similarity of Soukanzan (sou), Mangoku-ura (mng) and Akkeshi (akk) with introduced sites (Figure S5).

Finally, as briefly mentioned above, we found some evidence consistent with secondary introductions (i.e., movement of introduced thalli to other locations within the introduced range). The strongest evidence of secondary introduction came from the three sites from Long Island Sound and Narragansett Bay (nyc, lhp, ris). Bayesian clustering at *K*=5 (Figure 3, Figure S3) and the NJ analyses (Figure S5) suggested these sites were more closely aligned with WNA sites (elk, ens) than with other EUSA sites. In addition, blind assignment of introduced populations onto multivariate DAPC space created by genotypes from sites in the native range and two of the three introduced coastlines suggested introduced thalli may have more complex primary and secondary invasions (Figure S6). For example, EUSA and EU thalli were assigned to sites from the central Californian coast in addition to the northeastern coastline of Japan.

## DISCUSSION

Using a combination of mitochondrial and microsatellite genotyping, we discovered profound phylogeographic structure in the native range of *Gracilaria vermiculophylla*, reflecting low levels of contemporary gene flow. Introduced populations of North America and Europe exhibited mitochondrial and nuclear genotypes largely indistinguishable to those of the Pacific coastline of northeastern Japan (i.e., northern Honshu and southeastern Hokkaido Islands). Because this area historically served as the principal source for oyster exports worldwide, our results support anecdotal evidence that oysters were the key vector for the *G. vermiculophylla* invasion. We discuss each of these findings in detail below.

### Phylogeographic structure in the native range

In comparison to well-studied biogeographic provinces, such as those in northeast (NE) Atlantic, relatively little is known about the evolutionary history of the organisms that inhabit the northwest (NW) Pacific and it is not clear whether hypotheses drawn from patterns in other regions apply (Cheang *et al.* 2010). Phylogeographic studies in the NW Pacific have been biased towards commercially exploitable fish and mollusks and the majority have been underappreciated within English-language journals because of language barriers (Ni *et al.* 2014). Nevertheless, phylogeographic patterns have been found to be complex due to intricate topography and dynamic current systems (Wang 1999).

The marginal seas of South China Sea, the East China Sea and the Sea of Japan/East Sea may have served as glacial refugia, though intraspecific variation in demographic histories are common across taxa (Ni *et al.* 2014). He *et al.* (2015) found the amphibious mudskipper *Periophthalmus modestus* colonized northward from the South China Sea through the East China Sea along the coasts of Korea and Japan. Other studies have found higher haplotypic diversity in southern South Korea and southern Japan (i.e., East China Sea) as compared to Honshu and Hokkaido populations (Kim *et al.* 2012; Ho *et al.* 2014), suggesting northward colonization. However, though studies have found southern clade haplotypes decrease to the north (Azuma & Chiba 2015), no studies, to our knowledge, have found as sharp a break as observed in this study of *Gracilaria vermiculophylla* in the NW Pacific (Figure 1, Figure S1).

We found three shallow clades among mtDNA sequences, suggesting divergent demographic histories that were geographically correlated (Figure S1e). Two ‘C’ groups were found south of ˜35 °N, corresponding to warm temperate biogeographic provinces, whereas the ‘T’ group was found north of ˜35 °N corresponding to cold temperate biogeographic provinces (Spalding *et al.* 2007; Briggs & Bowen 2011). The ‘C’ clade with 10 haplotypes was found predominately in the East China and Yellow Seas, whereas, the other ‘C’ clade was found further east from South Korea to the Chiba Peninsula in Central Honshu (Figure S1). The ‘T’ haplotypes may have spread northward from the refugium in the Sea of Japan/East Sea, whereas, the ‘C’ haplotypes were restricted to warm-water provinces. Present-day maintenance of the ‘C’/’T’ boundary may reflect current patterns, such as the barrier at Cape Inubo where the warm-water Kuroshio Current meets the cold-water Oyashio current. It is difficult to assess which clade is most basal given the lack of sequence information for closely-related *Gracilaria* species.

Though the majority of studies have focused on broader scale patterns across the marginal seas of the NW Pacific from southern China to Japan (Ni *et al.* 2014), there is strong genetic differentiation between populations along the Sea of Japan/East Sea and the Pacific coast. For example, the eastern and western coastlines of Japan were genetically differentiated between populations of *Pterogobus* gobies (Akihito *et al.* 2008), the fucoids *Sargassum hemiphyllum* (Cheang *et al.* 2010) and *S. horneri* (Uwai *et al.* 2009) and the kelp *Undaria pinnatifida* (Uwai *et al.* 2006), but not in the gastropod *Littorina brevicula* (Azuma & Chiba 2015). The ‘T’ haplotype 6 was common on both coastlines of Japan at roughly 35 °N and the eastern coastlines of South Korea and Russia, but haplotypes 5 and 7 and 16, 18 and 28 were only sampled in the Sea of Japan/East Sea and Pacific Ocean populations, respectively (Figure S1a).

Microsatellite genotypes, on the other hand, were characterized by genetic differentiation that corresponded to a large degree with extant current patterns (Figure 3, Figure S3). For example, the southern Japanese sites, from southeastern Kyushu to the Chiba Peninsula and likely isolated by the Kuroshio Current, formed a large cluster with minimal admixture (Figure S3). Likewise, Sea of Japan/East Sea and Hokkaido sites formed a group that was consistent with the flow of the Tsushima Current. In contrast, the thalli sampled at Hayase River (hay) and Futatsuiwa (fut) that were genetically more similar to northeastern Honshu sites (mou and nag). It is possible that these reflect some movement of *Gracilaria vermiculophylla* thalli as a result of aquaculture practices (i.e., oysters or the alga itself). Finally, the similarity of thalli sampled at Shikanoshima (shk) to thalli sampled from source sites (akk, mng, sou) is likely due to the frequent fishery transport between Hokkaido and Kyushu (M. Nakaoka, *pers. comm.*), of which similar patterns have been found in the red seaweed *Gelidium elegans* (Kim *et al.* 2012).

*Pathway of the oyster? Gracilaria vermiculophylla* invaded low-energy, high salinity estuaries dominated by soft-sediments on all continental margins of North America (Bellorin *et al.* 2004, Kim *et al.* 2010), habitats in which oysters were seeded and cultivated in huge quantities. However, the first records of *G. vermiculophylla* along the WNA, EUSA and EU coastlines were 1979 (Bellorin *et al.* 2004), 1998 (Thomsen *et al.* 2006) and 1995 (Rueness 2005), respectively, which are decades after deliberate oyster movement (Barrett 1963; Ruesink *et al.* 2005), leading to speculation that other vectors (i.e., aquaculture, hull fouling or ballast water) were more likely (Ruiz *et al.* 2011), J. Carlton, *pers. comm*.).

Our results, however, point to oyster transport as the principal vector for the spread of *Gracilaria vermiculophylla*. Specifically, our genetic analyses assigned nearly all introduced algal thalli to three prominent sites of oyster aquaculture (reviewed in Barrett 1963, Byers 1999; Ruesink *et al.* 2005) along the northeastern coastline of Japan: Soukanzan (sou, Matsushima Bay), Mangoku-ura (mng, Mangoku Bay) and Akkeshi (akk, Lake Akkeshi). These support anecdotal evidence that the first oysters introduced to Puget Sound were from Akkeshi (Galtsoff 1932) and that subsequent introductions to California (Barrett 1963, Chew 1979) and France (Maurin & Le Dantec 1979, Goulletquer 1998) were mainly from Matsushima Bay (Chew 1979). While we cannot eliminate deliberate introductions for aquaculture, or incidental introductions from hull fouling or ballast water as playing a role, the simplest explanation is oyster introductions.

Surprisingly, and despite the fact that dozens of species were introduced worldwide with Japanese *Crossastrea gigas* from northeastern Japan (Byers 1999; Ruesink *et al.* 2005), we know only a single study in which this region was independently identified as an invasion source using genetic markers (i.e,. the mud snail *Batillaria attramentaria,* Miura *et al.* 2006). The dearth of studies that identified this historically important source region is due either to poor resolution of markers, poor sampling of native range populations or both. Our study and Miura *et al.* (2006) rigorously sampled the native and non-native ranges enabling the identification of intensive oyster aquaculture as the source populations, while simultaneously excluding other native sites as potential donor regions.

### Complexity of introduction history

There were likely multiple *Gracilaria vermiculophylla* invasions along each continental shoreline. Along the west coast of North America, estuaries in Tomales Bay (tmb), Elkhorn Slough (elk) and the Salish Sea (ptw) were composed of different genetic cohorts of *G. vermiculophylla.* This is consistent with the anecdotal histories of oyster introduction and cultivation in these locations (Barrett 1963; Byers 1999). Along the coastlines of northwestern Africa and Europe, we also found evidence of genetic structure consistent with multiple introductions. In Europe, there is a long and complicated history of oyster introductions that dates back to the 1600s directly from the northwest Pacific (Grizel and Héral 1991, O Foighil *et al.* 1998; Goulletquer 1998; Haydar & Wolff 2011, Lallias *et al.* 2015). In addition to *Crassostrea gigas* introductions, *C. virginica* was introduced from the EUSA to hatcheries and aquaculture facilities throughout Western Europe (Ruesink *et al.* 2005). Other invaders, such as *Sargassum muticum* and *Crepidula fornicata*, are thought to have been accidentally introduced to Europe as a result of *C. gigas* oyster introductions from Japan or British Columbia (Farnham *et al.* 1973) or *C. viriginica* imported from EUSA (Riquet *et al.* 2013), respectively. The similarity of European sites to some WNA and EUSA sites suggests that the invasion of the European coastline may reflect primary invasions from Japan and secondary invasions from WNA, EUSA or both. With our current set of microsatellite genotypes, we cannot at present distinguish between these two alternatives.

Along the east coast of the United States, there were at least two different invasions; one to Long Island Sound (nyc, lhp) and Narragansett Bay (ris) and the other to the outer coast of Virginia and the Chesapeake Bay (Figure 3). The first introduction stayed localized to Long Island Sound to the Cape Cod Peninsula and maintains a genetic constitution that is more similar to the EU and WNA than to the other populations along the EUSA. The second introduction rapidly spread along the eastern seaboard to New Hampshire (grb) and south to the Carolinas and Georgia and may have been introduced directly from Japan (Mann 1979). However, as with the EU, we cannot distinguish among primary and secondary invasions with our current data.

Though the first records of EUSA *Gracilaria vermiculophylla* only date back to 1998, this alga likely occurred in the Chesapeake Bay and along New England coastlines for longer than currently thought and was misidentified as the native congeneric *G. tikvahiae* (Thomsen *et al.* 2006; Nettleton *et al.* 2013). *Crassostrea gigas* was introduced to Maine and Massachusetts in 1949 and the 1970s (Dean 1979, Hickey 1979), Connecticut in the 1940s (Loosanoff & Davis 1963), New Jersey in the 1930s, Delaware in 1962 and Maryland in the 1970s (Andrews 1980). Likewise, the arrival of an oyster pathogen in the Chesapeake Bay was linked to rogue plantings of *C. gigas* from Japan, California, the northwest Pacific or some combination of these oyster cultivation regions (Burreson *et al.* 2000). Taken together, our contemporary genetic data strongly point to the movement of oysters as the main vector and pathway for moving *Gracilaria vermiculophylla* across oceanic basins.

### Secondary spread within coastlines

Our genetic data suggest different patterns of secondary spread once *Gracilaria vermiculophylla* invaded each of the continental margins of the Northern Hemisphere. Along the WNA coastline, the genetic structure of *G. vermiculophylla* was the most profound, relative to other continental margins. This suggests that multiple primary invasions arrived from Japan and that secondary spread was likely less important than along the EUSA and EU coastlines. This is a tentative conclusion, however, as the geographic extent of the introduction of *G. vermiculophylla* is poorly characterized along the WNA coast. In 2015, for example, we found *G. vermiculophylla* in Bodega Bay (bob), Tomales Bay (tmb) and San Diego (E.E. Sotka and S.A. Krueger-Hadfield, *unpubl. data*). Grosholz and Ruiz (2009) documented the dramatic change in abundance of what they called *Gracilariopsis sjoestedtii* in Bodega Harbor as a result of the invasion of the European crab *Carcinus maenas*. They analyzed aerial photographs, but performed their study in the same area we sampled *G. vermiculophylla* in this study. An herbarium sample at the Bodega Marine Lab from the late 1960s from this same area was labeled *Gracilaria* cf. *vermiculophylla* (P.G. Connors, *pers. comm.*) and has subsequently been identified as *G. vermiculophylla* (S.A. Krueger-Hadfield and K.A. Miller, *unpubl. data*).

Relative to the case along the WNA coastline, the magnitude of secondary spread along the EU and EUSA coastlines is less speculative. Along the EUSA, the high levels of diversity observed in the sites studied along the eastern shore of Virginia (Figure 4; see also Gulbransen *et al.* 2012) strongly suggest that the outer coast of Virginia and the Chesapeake Bay were the initial sites of introduction for the larger EUSA invasion. Rapid colonization southward has been documented, such as throughout the estuaries of South Carolina (Byers *et al.* 2012), possibly facilitated by the Inter-Coastal Waterway (Freshwater *et al.* 2006). Consistent with this southward invasion pathway, there was a decline in allelic diversity from Virginia to Georgia (Figure 4).

Along the coastlines of the British Isles, Europe and northern Africa, our data are consistent with multiple invasion events into France and Portugal, followed by secondary spread to other estuaries. In the Marennes-Oléron basin, for example, *G. vermiculophylla* thalli from Loix (frl) and Marennes (fme) belonged to different genetic cohorts (Figure S3). This basin is an area of intense aquaculture and the similarity of the sites sampled in Brittany (fdm, pol, fpa) to the thalli sampled at Marennes (fme) may be the result of secondary spread as oysters are commonly moved throughout French cultivation facilities in the English Channel, Bay of Biscay and Mediterranean throughout their life cycle (Goulletquer 1998). By contrast, sites sampled along the Jutland Peninsula (hei, dhd, nib, nor, man) were largely composed of a single genetic cluster from *K* = 2 to *K* = 23 (Figure 3, Figure S3) and exhibited lower estimates of diversity than other European sites (Figure 4). Gracilarioid seaweeds were not found in the Kiel fjord, for example, until the mid-2000s, suggesting the invasion of Jutland estuaries is more recent (F. Weinberger, *pers. obs.*). These sites also exhibited genetic signatures of extensive vegetative fragmentation where we previously found few unique genotypes (Krueger-Hadfield *et al.* 2016). On the other hand, oysters have repeatedly been introduced into the German Wadden Sea from Portugal (Mayer-Waarden 1964), which was reflected in the genetic similarity of of the Jutland populations with those of the Ria d’Aveiro in Portugal (Gafanha/gaf and Estrareja/est; Figure S5a) We did not sample Mediterranean populations, which may help to identify pathways that are cryptic to this current study, such as the introduction of the Manila clam (Sfriso *et al.* 2010).

The vector of secondary spread along the EU and EUSA coastlines is uncertain. Spread of *Crassostrea gigas* throughout the East Frisian Wadden Sea may be responsible for the encroachment of *Gracilaria vermiculophylla* along the Jutland Peninsula (Schmit *et al.* 2008 *Helgol Mar Res*). Indeed, *G. vermiculophylla* is often found near oyster farms (Rueness 2005). Spread of *G. vermiculophylla* along the EUSA may be via shrimp nets and crab pots, as *G. vermiculophylla* is known to be a nuisance in some estuaries to shrimpers (Wilson Freshwater *et al.* 2006). Nyberg and Wallentinus (2009) suggested migrating seabirds may also be responsible for the transport of small fragments of *G. vermiculophylla*. Seabird sanctuaries in closed lagoons along the German Baltic Sea coast that are inaccessible to aquaculture, boat traffic and other human activities often harbor isolated populations of *G. vermiculophylla* (F. Weinberger and M. Hammann, *pers. obs.*). Birds have also been known to use gracilarioid seaweeds in nests (Arnold 1968), including *G. vermiculophylla* (S.A. Krueger-Hadfield and S.J. Shainker, *pers. obs.*).

### Conclusions

*Gracilaria vermiculophylla* is now recognized as one of the most widespread and abundant marine invaders in the Northern Hemisphere (Saunders 2009; Kim *et al.* 2010; Krueger-Hadfield *et al.* 2016) and has transformed the ecosystems to which it has been introduced (Thomsen *et al.* 2009; Byers *et al.* 2012). Nevertheless, this cryptic invader has been lurking in estuaries throughout the Northern Hemisphere for decades without recognition. This may reflect the fact that phycologists are more likely to study rocky shores than soft-sediment estuaries where *G. vermiculophylla* has largely invaded, or the fact that cryptic species are common in seaweeds (Krueger-Hadfield et al. in review). An analogous example comes from Provan *et al.* (2007), who used herbarium specimens coupled with molecular markers to conclude that the invasive strain of *Codium fragile* spp. *tomentosoides* colonized sites around the world for at least 100 years longer than previously reported from contemporary records.

Our identification of a source region has implications for understanding the evolutionary ecology of the *Gracilaria vermiculophylla* invasion, in particular. Previous studies have suggested introduced populations evolved greater tolerance for heat stress and resistance to herbivory and bacteria during the invasion (Hammann *et al.* 2013a; 2016). However, these studies compared introduced populations with Korean and Chinese native populations that do not represent the source of the invasion. If there is geographic variation in the native range in these traits, then it is possible that the source populations had already evolved these traits before the invasion (Bossdorf *et al.* 2008; Estoup & Guillemaud 2010; Colautti & Lau 2015).

More broadly, Carlton (2009) argued convincingly that fewer assumptions should be made about the status of a given taxon (e.g., native or cryptogenic) without appropriate morphological and molecular analyses. This is particularly true for understudied groups in under-explored habitats. We extend Carlton’s (2009) suggestion to not only thoroughly explore the systematics of invaders, but also invasion vectors and pathways. Fewer assumptions should be made about the likelihood of a given vector without appropriate historical and contemporary genetic sampling and analyses.

## ACKNOWLEDGEMENTS

Microsatellite primer sequences were deposited in GENBANK, accession numbers KT232089-KT232097 and KT232099 (Kollars *et al.* 2015). Microsatellite genotypic data were deposited in DRYAD, entry *TBA*. Mitochondrial sequence data were deposited in GENBANK (Acc. No. *TBA*).

We are grateful for everyone who provided algal samples (see Table S1 for complete list), and in particular, R. Hadfield, C. Sotka, M. Nakaoka, M. Kamiya and H. Endo. Thank you to J. Carlton, M. Valero and C. Destombe for insightful discussions; B. Flanagan for help with DNA extractions; K. Hill-Spanik for help with phylogenetic analyses; K. Holcombe at the Chincoteague National Wildlife Refuge (FWS Special Use Permit SUP 51570-2014-013) and B. Hughes (Elkhorn Slough National Estuarine Research Reserve) for site access; and G. Saunders for field locations in British Columbia. This project was supported by NSF BIO-OCE-1057713, BIO-OCE-1057707, BIO-OCE-1357386; a College of Charleston Graduate Research Grant; the Phycological Society of America Grants-in-Aid-of-Research; Zostera Experimental Network Graduate Research Fellowship (NSF OCE-1031061); and LLUR-Schleswig-Holstein.

The scientific results and conclusions, as well as any opinions expressed herein, are those of the author(s) and do not necessarily reflect the views of NOAA or the Department of Commerce. The mention of any commercial product is not meant as an endorsement by the Agency or Department.

## AUTHOR CONTRIBUTIONS

SAKH, NMK and EES conceived the study; SAKH, NMK, JEB, MH, SJS, RT, FW and EES collected samples; SAKH, NMK and SJS extracted DNA; SAKH, NMK, SJS, TWG and DM generated data; SAKH, AES and EES analyzed data; JEB and AES contributed to discussions; SAKH and EES wrote the manuscript; all authors approved the final manuscript.

